# Elucidation of the second-phase freezing acclimation mechanism: plants accumulate sugars via photosynthesis under frozen conditions

**DOI:** 10.64898/2026.10.14.756911

**Authors:** Kenji Sugita, Matsuo Uemura, Atsushi J. Nagano, Manabu Watanabe, Yukio Kawamura

## Abstract

Overwintering plants increase their tolerance to freezing in two phases: the first-phase, known as cold acclimation (CA), occurs at chilling temperatures, followed by second-phase acclimation at sub-zero temperatures. We previously showed that second-phase freezing acclimation (2PFA), in which cold-acclimated *Arabidopsis* is exposed to -2°C with extracellular ice formation under light, confers the highest freezing tolerance (LT_50_ below -20°C) through a pathway requiring photosynthetic electron transport, possibly independent of the CBF-COR pathway. However, how light enhances freezing tolerance in frozen plants remains unclear. Here, we developed a system to measure chlorophyll fluorescence in intact plants maintained in a frozen state and found that PSII retained substantial photochemical capacity (*Fv/Fm*) in frozen leaves. Comparative transcriptomics revealed that 2PFA under light induced genes involved in starch, sucrose and galactose metabolism. Consistently, soluble sugars, mainly sucrose, glucose and fructose, accumulated to levels approximately 8-fold higher than those in CA plants, and total sugar content was strongly correlated with freezing tolerance. This accumulation was suppressed by the photosynthetic electron transport inhibitor DCMU. These results suggest that photosynthetic activity is maintained in frozen plants and supplies sugars that may contribute to the freezing tolerance acquired during second-phase acclimation, beyond the level achieved by first-phase CA, unveiling a previously unrecognized mechanism in the winter survival strategy of plants.

## 1 Introduction

Plants that grow in temperate and frigid zones are exposed to subzero temperatures during winter, and they develop freezing tolerance at the beginning of fall in a stepwise manner. A first-phase process toward overwintering from late fall proceeds at non-freezing temperatures, and plants increase their freezing tolerance, which is known as cold acclimation (CA). A second-phase process proceeds after mid-winter, in which plants experience temperatures below 0°C, and freezing tolerance is further enhanced. This phenomenon is known as second-phase cold hardening (2PH) or subzero acclimation. Recently, our studies have shown that the 2PH process can be divided into four patterns that is plants are exposed to subzero temperatures with (second phase freezing acclimation; 2PFA) or without ice (second phase supercooling acclimation; 2PSA) under dark or light conditions (Sugita et al., 2024).

The 2PH process is a general phenomenon in overwintering plants, and freezing is a key signal for significant enhancement of freezing tolerance. In particular, 2PFA under light conditions was reported to lead to the highest freezing tolerance among the 2PH patterns, with an LT_50_ value of approximately below -20°C in *Arabidopsis* (Sugita et al., 2024). Despite the fact that there have been very few reports, studies on 2PH in the last quarter-century have focused on herbaceous plants such as *Arabidopsis*, wheat, and oat (Livingston & Henson, 1998; Herman et al., 2006; Livingston et al., 2007; Le et al., 2008, 2015; Takahashi et al., 2019, 2021, Sasaki et al., 2026). In woody plants, subzero temperatures or frost conditions can cause a second stage of CA, which is known to improve freezing tolerance (a 2PH-like phenomenon) (Weiser, 1970; Sakai & Larcher, 1987). Thus, 2PH is a universal phenomenon in overwintering plants, and a detailed analysis of the physiological effects of combined freezing and light conditions on plants during the 2PH process is important to understanding plant overwintering phenomena.

Studies on CA processes have shown that light is an important factor in enhancing freezing tolerance. For example, sugars such as sucrose, glucose, fructose, and raffinose accumulate in many plants, including monocotyledonous, dicotyledonous, herbaceous, and woody plants during CA under light conditions (Guy et al., 1992; Sasaki et al., 1996; Sauter et al., 1996; Savitch et al., 2000; Kamata and Uemura 2004; Klotke et al., 2004; Lee et al., 2013; Juurakko et al., 2021), and several studies have reported their association with photosynthesis (Guy et al., 1992; Wanner & Junttila, 1999; Savitch et al., 2000). An increase in sugar content is positively correlated with enhanced freezing tolerance (Sakai & Larcher, 1987; Strand et al., 2003; Yano et al., 2005; Lee et al., 2013), suggesting that a phenomenon presumed to be involved in tolerance to freeze-dehydration. In freeze-tolerant plants, ice crystals, which are always extracellular, are first formed, and because water molecules in the crystalline state have lower chemical potential than those in the liquid state, extracellular freezing results in the movement of water from the intracellular space to the extracellular space. Consequently, plant cells experience dehydration stress under freezing conditions. Electron microscopic observations of plant cells under freezing conditions revealed that certain abnormal structures exist in the plasma membrane at temperatures that cause irreversible injury to cells (Gordon-Kamm & Steponkus, 1984; Steponkus et al., 1993; Uemura et al., 1995; Fujikawa et al., 1999; Nagao et al., 2008). These structures are thought to be caused by the abnormal proximity of the plasma membrane to the intracellular membrane or salt stress from freeze-dehydration. Sugars may act as a direct buffer to prevent the fusion of cellular membranes induced by freeze-dehydration (Steponkus, 1984; Uemura et al. 2003). The *Arabidopsis* mutant *sfr4*, which does not show improved freezing tolerance to CA, is primarily caused by lack of sugar accumulation during CA (McKown et al. 1996; Warren et al. 1996; Uemura et al. 2003). Therefore, sugar accumulation may play an important role in enhancing freezing tolerance.

Several sugar synthesis genes, particularly with regard to sucrose synthesis, and transcripts that encode *Sucrose-phosphate-synthase (SPS)* and *Sucrose synthase (SuSy)* have been reported to be upregulated in response to low temperatures (Strand et al., 1997; Gilmour et al., 2000; Fowler & Thomashow, 2002; Kaplan et al., 2007). C-repeat binding factors/dehydration-responsive element binding factors (CBFs/DREBs, a type of transcription factor) expressed in the early phase of CA have been shown to be central factors in the regulation of acclimation processes (Thomashow, 1999, 2001; Park et al., 2015; Jia et al., 2016; Zhao et al., 2016; Song et al., 2021). In *Arabidopsis*, overexpression of the *CBF3* gene does not affect the expressions of *SPS* and *SuSy* but contributes to an increase in sugars during CA (Gilmour et al., 2000). RNA sequencing (RNA-seq) analysis confirmed that the expressions of *SPS* and *SuSy* were suppressed in *cbfs* triple-deficient *Arabidopsis* compared to that in the wild type, but they increased during acclimation (Jia et al., 2016; Zhao et al., 2016). These results indicate that the increase in sugars during CA may not only be due to the CBF pathway but also due to other pathways.

Our previous study showed that the 2PFA process under light conditions resulted in the maximum freezing tolerance of plants via chloroplast electron transfer signals, but the transient increase in *CBF* genes during this process was remarkably low compared to 2PH under the dark (Sugita et al., 2024). These results suggest that the main pathway involving photosynthesis other than CBF probably enhanced freezing tolerance in the 2PFA process under light conditions. The big question remains is “Do frozen plants actually undergo photosynthesis?”. Although photosynthesis has been observed in Antarctic lichens under freezing conditions (Barták et al., 2007; Andrzejowska et al., 2024), there have been no studies on higher plants, and there are some suspicious presuppositions regarding photosynthesis under freezing conditions (Öquist & Huner, 2003).

This study aimed to elucidate the mechanism of enhanced freezing tolerance in *Arabidopsis* in the 2PFA process under light conditions. We first constructed an experimental system that enabled chlorophyll fluorescence spectroscopy under freezing conditions and measured the photosynthetic activity. Additionally, we conducted RNA-seq analysis and found that changes in sugar metabolism were key pathways in 2PFA plants under light conditions. Finally, we examined sugar accumulation during the 2PFA process under light conditions and its relationship with photosynthesis. Our results showed that a significant amount of sugar accumulated during the 2PFA process via photosynthesis and gene expression even when plants were frozen, suggesting that significant freezing tolerance may be related to these processes

## 2 Materials and Methods

### 2.1 Plant materials and growth conditions

Seeds of *Arabidopsis thaliana* ecotype Columbia (Col-0) were surface-sterilized with 20% (v/v) bleach and 1% (v/v) sodium dodecyl sulfate and placed on Hoagland’s medium containing 1% (w/v) sucrose and 1% (w/v) agar in square (96 mm ×96 mm) Petri plates. After two days of stratification at 4°C in the dark, plants were grown in a controlled-environment chamber at 23°C for 2 wks under a 16-h light/8-h dark photoperiod with light supplied at 80 μmol m^-2^ s^-1^ by fluorescent lamps. These plants were designated as non-acclimated (NA) plants. The NA plants were transferred into a growth chamber at 2°C under a 12-h light/12-h dark photoperiod with light supplied at 80 μmol m^-2^ s^-1^ by fluorescent lamps and grown for 7 d as the CA treatment. The CA plants were then transferred to a freezing chamber that could maintain stable subzero temperatures even under light conditions. This was achieved by adding circulation fans and LED lights to a ready-made freezing chamber (IN602N, Yamato Scientific, Tokyo, Japan) as described in a previous study (Sugita et al., 2024). This chamber was used to incubate the plants for three days at -2°C either in the dark or under a 12-h light/12-h dark with light supplied at 80 μmol m^-2^ s^-1^ as the second-phase freezing acclimation (2PFA) treatment. The plates were kept at -2°C for 10 min in the freezing chamber, and ice was induced to form by momentarily touching the agar with liquid nitrogen. In the second-phase supercooling (2PSA) treatment, CA plants were transferred to the freezing chamber at -2°C in the same way as in the 2PFA treatment, but ice was not induced to form on the agar. As one of the controls, CA plants were subjected to a further CA treatment (CA+) under the same conditions as the 2P treatment at 2°C to investigate a possibility that effects other than temperature and freezing might affect the result.

All experiments were repeated at least three times, and mean values were calculated.

### 2.2 Measurement of sugar content

The aerial parts of each sample (∼0.1 g) were frozen in liquid nitrogen. The plant materials were homogenized with 80% (v/v) ethanol using a pestle. The homogenate was boiled for 30 min on a heat block at 80°C and then centrifuged for 5 min. The residues were rehomogenized with 80% (v/v) ethanol and centrifuged as described above. These extraction steps were repeated three times. The supernatants were dried on a heat block at 80°C. Leaf pigments were removed using chloroform, methanol, and water (chloroform:methanol: water, 1:1:0.9, v/v/v). The extracts were dried on the heat block at 80°C and dissolved in 0.5 mL of distilled water and then filtered through a 0.22 μm pore filter membrane. Sucrose, raffinose, glucose, and fructose were analyzed using high-performance liquid chromatography (HPLC) using SUGAR SC1011 (Resonac, Tokyo, Japan) and SUGAR KS-801 (Resonac) columns at 30°C with a refraction index detector. Samples (5μl) were injected and eluted with ultrapure water at a flow rate of 0.2 ml min^-1^. Each sugar was quantified based on a calibration curve using an external standard method. For measurement of trehalose content, we used NH2P-50 4E (Resonac) columns at 35°C with a refraction index detector, and samples (10μl) were injected and then eluted with 75% (v/v) acetonitrile at a flow rate of 0.8 ml min^-1^.

### 2.3 Freezing tolerance test

Seedlings grown on agar plates were placed vertically in the dark in an SU-641 controlled-temperature chamber (Espec, Osaka, Japan) and rotated at 1 rpm to equalize all plate temperatures through air circulation using a fan. After 10 min at -2°C, for the 2PSA plants, ice crystals were formed on the agar by touching a corner of the plant with liquid nitrogen for 3 seconds. For 2PFA plants, this treatment was not applied since ice crystals had already been formed. The plates were kept for 2 h at -2°C to freeze the agar and plants completely, and the temperature was gradually lowered to -20°C at -2°C h^-1^ in the dark. After removing the plates at the specified temperature, plants were transferred to a growth chamber at 23°C with a 16-h light/8-h dark photoperiod with light supplied at 80 μmol m^-2^ s^-1^ by fluorescent lamps. The plants were kept under these conditions for 1 d to allow thawing, and the survival rate was then measured.

### 2.4 Measurement of survival rate

Survival rate was measured using photosystem II (PSII) (F_v_/F_m_) on intact plants as described in previous study (Hiraki et al., 2020; Sugita et al., 2024). Briefly, the chlorophyll fluorescence of dark-adapted *Arabidopsis* plants (15 min) was measured using a Fluor Cam instrument (800 MF; Photon Systems Instruments, Brno, Czech Republic). The survival rates were calculated as the ratio of the maximum quantum yield before and after freezing multiplied by the ratio of the area showing detectable fluorescence before and after treatment. For details on the calculation, see Hiraki et al. (2020) and Sugita et al. (2024). The LT_50_ was calculated via linear regression using data from the two temperatures at the interval between less than and more than 50% damage.

### 2.5 Treatment with a photosynthetic electron transport inhibitor

CA seedlings were misted twice with 0.5% (v/v) ethanol with or without 50 μM 3-(3,4-dichlorophenyl)-1,1-dimethylurea (DCMU). The experiment was performed at 2℃ under green filtered low light to prevent photosynthesis. After 1 h at 2℃ under dark conditions, the plants were transferred into the growth chamber at -2°C with ice crystallization for 3 d under dark conditions or a 12-h light/12-h dark photoperiod with light supplied at 80 μmol m^-2^ s^-1^ by fluorescent lamps. The sugar contents of the aerial parts of the plants were then measured.

### 2.6 Chlorophyll fluorescence measurement

To measure the chlorophyll fluorescence of plants under freezing conditions, *Arabidopsis* seedlings were placed on Hoagland medium containing 1%(w/v) sucrose and 1%(w/v) agar in a Petri dish (diameter: 40 mm). After stratification, the plants were grown at 23°C for 2 wks and transferred into the growth chamber at 2°C for 7 d to induce cold acclamation. The plants were then placed on the growth chamber at -2°C, and ice crystallization was inoculated to agar by momentarily touching the plate with liquid nitrogen after 10 min of incubation. The plants were kept at -2°C under dark conditions or 12-h photoperiod at 80 μmol m^-2^ s^-1^.

To measure the chlorophyll fluorescence in the frozen state, a low-temperature water tank (Tokyo Rikakikai, Tokyo, Japan) was used to circulate antifreeze (EtOH) through a jacketed beaker (**Fig. 1**). Plates after incubation at -2°C for 0, 24, 48, and 72h were placed in the jacketed beaker. The chlorophyll fluorescence was measured while the agar was maintained at freezing temperature. The insulating sheets were wrapped around the jacked beaker and the hose was connected to a low-temperature bath to stabilize the temperature (**Fig. 1**).

**Fig. 1.**
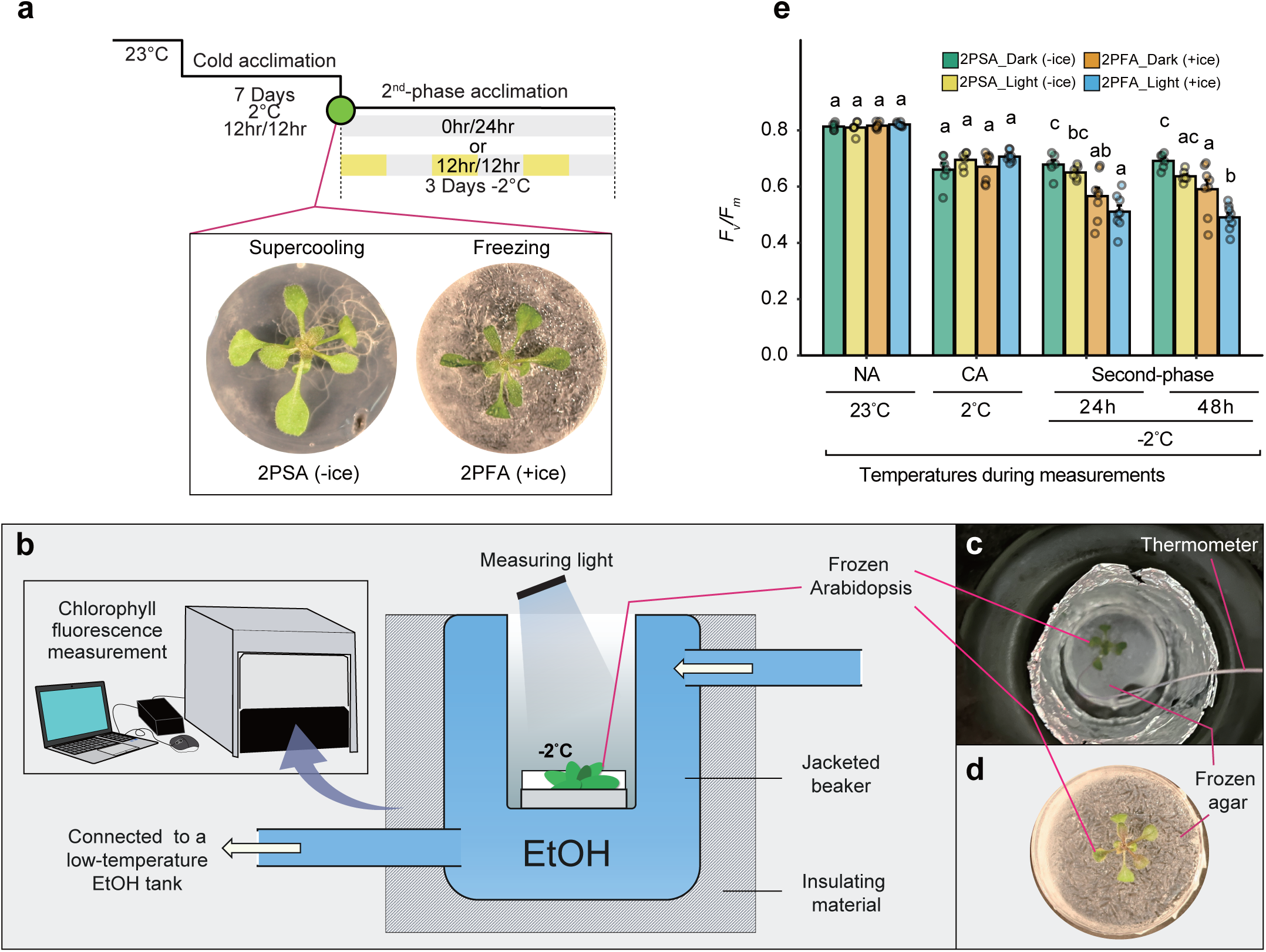
*Fv/Fm* of plants during 2^nd^-phase freezing acclimation (2PFA) and 2^nd^-phase supercooling acclimation (2PSA) in the dark and light conditions after CA. (a) Schematic diagram of the experiment for measurements of chlorophyll fluorescence. Antifreeze solutions (EtOH) were circulated in a jacked beaker using a low-temperature water tank. Insulating sheets were wrapped around the beaker and hose connecting between the beaker and low-temperature bath. (b) Plants were placed in the beaker, and the leaf temperature was kept at -2°C. (c) Ice was inoculated to agar by touching the plates with liquid nitrogen after 10 minutes of incubation at -2°C. (d) *Fv/Fm* was measured in the plants that were freezing or supercooling acclimated at -2°C in the dark or light conditions at 0, 24, and 48 h. As a control, Fv/Fm of the same samples used for the 2PFA in the dark and light conditions were measured when the plants were grown at 23°C for 2 weeks (NA). Error bars indicate SE (*n*=3-8). Different letters at each temperature show significant difference at *p* < 0.01 (Tukey-Kramer test).

### 2.7 RNA extraction

The samples of CA, CA+, 2PFA treatment, and 2PSA were collected at 0, 24, and 48 h for RNA-seq. RNA was extracted from the aerial parts of each sample using an ISOSPIN Plant RNA kit (Nippon Gene, Tokyo, Japan) according to the manufacturer’s instructions.

### 2.8 RNA-Seq analyses

Total RNA (500 ng) was used as the input of each sample for library preparation. Lasy-Seq was used to prepare the RNA sequencing (RNA-seq) library (Kamitani et al., (2019)) version 1.1 (https://sites.google.com/view/lasy-seq/). The library was sequenced using HiSeqX (Illumina, San Diego, CA, USA) with paired-end 100+100 bp mode. All obtained reads were first subjected to quality trimming using Trimmomatic to remove adapter sequences and low-quality bases. Subsequently, the trimmed reads were mapped on each reference and quantified using RSEM (version 1.3.3) and Bowtie (version 1.2.3) in Docker, dynacomkobe/biodocker_rnaseq_pipeline: ver.0.5.0 to convert raw data obtained via sequencing into count data. The Trimmed Mean of M values (TMM) method implemented in edgeR was used to normalize the RNA-seq data and perform the likelihood ratio test (LRT). Furthermore, the *k*-means method, Gene Ontology (GO) analysis, and Kyoto Encyclopedia of Genes and Genomes (KEGG) pathway gene set enrichment analyses were performed with R (version 4.3.2) software using Fragments Per Kilobase of exon per million mapped fragments (FPKM), which is an expression level normalization method. Transcriptome sequences are available in the DDBJ Sequence Read Archive repository (https://www.ddbj.nig.ac.jp/index-e.html) under accession numbers DRR793933-DRR793992.

### 2.9 Statistical analysis

Statistical significance of differences between two plant groups was determined using Student’s *t*-test (*p* < 0.05) in Excel (version 16.83) and one-way ANOVA with the Tukey-Kramer test for multiple comparisons (*p* < 0.01) using R (version 4.3.1) in RStudio (2025.05.1+513).

## 3 Results

### 3.1 Chloroplastic electron transfer during the 2^nd^-phase process under light conditions

Previous studies by our group showed that the CA plants treated with freezing (2^nd^-phase freezing acclimation: 2PFA) or without freezing (2^nd^-phase supercooling acclimation: 2PSA) on the agar at -2°C under light conditions notably had enhanced freezing tolerance (Sugita et al., 2024). Notably, 2PFA plants (in a frozen state during 2^nd^-phase acclimation) exhibit significantly higher freezing tolerance than 2PSA plants (in a supercooled state during 2^nd^-phase acclimation) at the same temperature of -2°C (Sugita et al., 2024; **Fig. 1a**). The increase in freezing tolerance of 2PFA plants was decreased by 3-(3,4-dichlorophenyl)-1,1-dimethylurea (DCMU), suggesting that the photosynthetic electron transport pathway is involved in the 2PFA process under light conditions (Sugita et al., 2024). Thus, to gain further insight into chloroplastic electron transfer during the 2PFA process, we evaluated the function of Photosystem II (PSII) of plants under freezing conditions (**Fig. 1**). We constructed an experimental system using a chlorophyll fluorescence spectrometer that could measure photosynthetic activity in intact (non-destructive) frozen plants (**Fig. 1b-d**). First, we analyzed the effect of PSⅡ photochemical efficiency on the chlorophyll fluorescence *Fv/Fm* ratio at cold and freezing temperatures (**Fig. 1e**). CA plants were subjected to 2PFA treatment for 24 and 48 h at -2°C under dark conditions or light conditions. The *Fv/Fm* values of NA, CA, and 2PSA plants were approximately 0.8 and 0.7, respectively. The *Fv/Fm* value of the 2PFA plants at 24 and 48 h decreased to ∼0.5. The difference between the values of the 2PFA plants under light and dark conditions was little. The decreased *Fv/Fm* values in 2PFA plants suggest that the photosynthetic activity in the electron transport system in chloroplast is reduced.

These results suggest that the electron transport system in chloroplast remains active during the 2PFA and 2PSA process, even though the PSⅡ photochemical efficiency is suppressed.

### 3.2 Transcriptional profiling reveals a unique gene cluster responsive to freezing under light conditions

We performed a large-scale gene expression analysis using RNA sequencing (RNA-seq) to understand the molecular mechanism by which 2PFA plants enhance their freezing tolerance under light conditions at the transcriptional level. In this study, CA plants were treated with freezing (2PFA) or without freezing (2PSA) on the agar at -2°C either in the dark or under a 12-h light/12-h dark photoperiod (**Fig. 2a**). As one of the controls, CA plants were subjected to a further CA treatment (CA+) under the same time conditions as the 2P treatment at 2°C to investigate a possibility that effects other than temperature and freezing might affect the result. Cluster analysis was performed to analyze the gene expression trends in the plants under various treatments (CA, CA+, 2PSA, and 2PFA). The genes were classified into 12 clusters (A-L) using the *k*-means method (**Fig. 2b**). In clusters A (*N*=473), H (*N*=252), and K (*N*=666), the gene expression levels were upregulated only in the CA+ plants, 2PSA plants under light conditions, and 2PFA plants under dark conditions. In clusters E (*N*=1766) and F (*N*=475), the gene expression was upregulated only in CA plants. In clusters B (*N*=175) and C (*N*=205), the gene expression levels were upregulated in CA and CA+ plants and 2PFA plants under the dark conditions. In cluster G (*N*=155), the gene expression in CA and 2PSA plants under light conditions and 2PFA plants either under dark or light conditions were upregulated, and those in CA+ and 2PSA under dark conditions were downregulated. Genes classified in cluster I (*N*=197) showed upregulated expression levels in the 2PFA plant under light conditions compared with those in the other plants in this study. In cluster J (*N*=252), the gene expression levels in the 2PFA and 2PSA plants under dark conditions were upregulated, whereas those in the CA, CA+, 2PSA, and 2PFA plants under light conditions were downregulated. In cluster L (*N*=267), the gene expression level in 2PSA plants under light conditions and 2PFA plants under dark or light conditions were upregulated, whereas the gene expression levels in CA and CA+ were downregulated. We focused on the cluster I genes that showed upregulated expression levels in the 2PFA plants under light conditions (**Fig. 2c**) to elucidate the process by which these plants achieve their maximum freezing resistance.

**Fig. 2.**
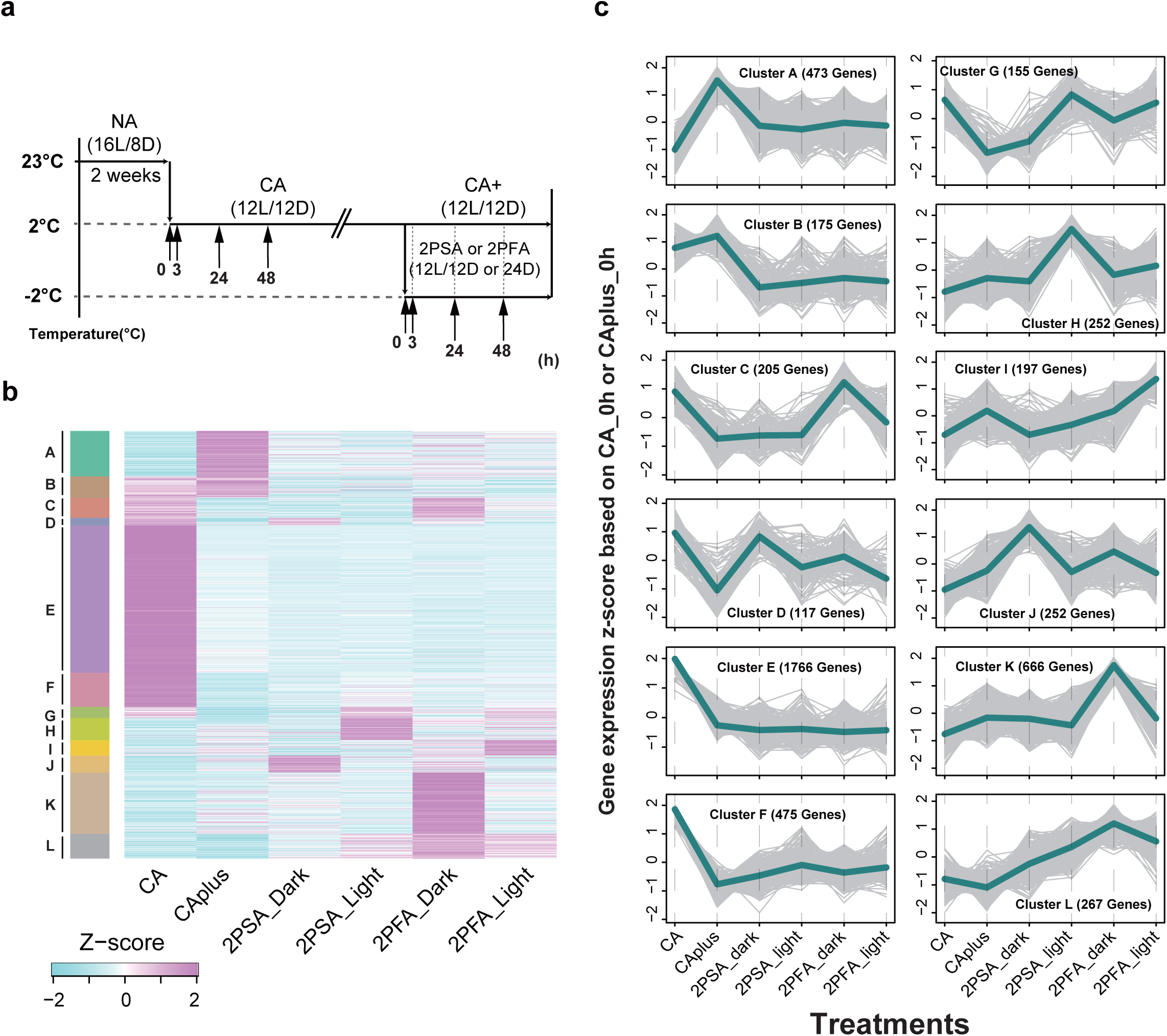
Genes from all treatment groups (CA, CA+, 2PSA, and 2PFA) into 12 clusters (A-L) using the *k*-means method. (a) Overview of experimental design. RNA was extracted from plants at 0, 3, 24, and 48 h after transfer into a growth chamber at 2°C from 23°C (CA samples). In addition, RNA samples were extracted at 0, 3, 24, and 48 h after transferring CA7 plants (cold-acclimated for 7 days at 2°C) into a freezing chamber at -2°C with or without ice crystallization (2PFA and 2PSA, respectively) in the dark or light conditions, or after keeping CA plants for a further 7 days at 2°C (CA+). The black arrow indicates the timepoints of the sampling. (b) Heatmap analysis of genes responding to CA, CA+, 2PSA, and 2PFA under dark or light conditions. The genes were classified into 12 clusters (Cluster A-L) using the *k*-means method, and random 1000 genes of them were visualized in the heatmap. The *k*-means analysis was performed with R (version 4.3.2) using RStudio (2025.05.1+513). (c) Line graphs showing the kinetics of changes in gene expression profile in all clusters shown in (b). The thin gray lines represent the individual gene expression trajectories of the genes within the cluster (Cluster A-L). A bold line represents the mean trajectory for the cluster.

### 3.3 Time-dependent transcriptional regulation only in the 2PFA plants under light conditions

A heat map of the time series analysis from 3 to 48 h in the CA, CA+, 2PSA, and 2PFA plant groups was constructed to examine the specific expression variations of genes belonging to Cluster I in detail (**Fig. 3a**). The results showed that the gene expression in 2PFA plants under light conditions was upregulated at 3, 24, and 48 h, and the expression patterns varied at each time point. Therefore, we used the *k*-means method to classify the expression patterns according to time. The 197 genes in the 2PFA plants under light conditions at 3, 24, and 48 h were further classified into four clusters (1-4) (**Fig. 3b**). The expression levels of the genes in cluster I-1 (*N* =53) rapidly increased within 3 h and then decreased at 24 h. In cluster I-2 (*N* =35), the expression levels of the genes increased at 3 and 24 h and decreased at 48 h. The gene expression levels in cluster I-3 (*N*=47) were slowly upregulated within 3 h, rapidly upregulated at 24 h, and then slowly downregulated at 48 h. In cluster I-4 (*N* =62), the gene expression levels were upregulated at 48 h.

**Fig. 3.**
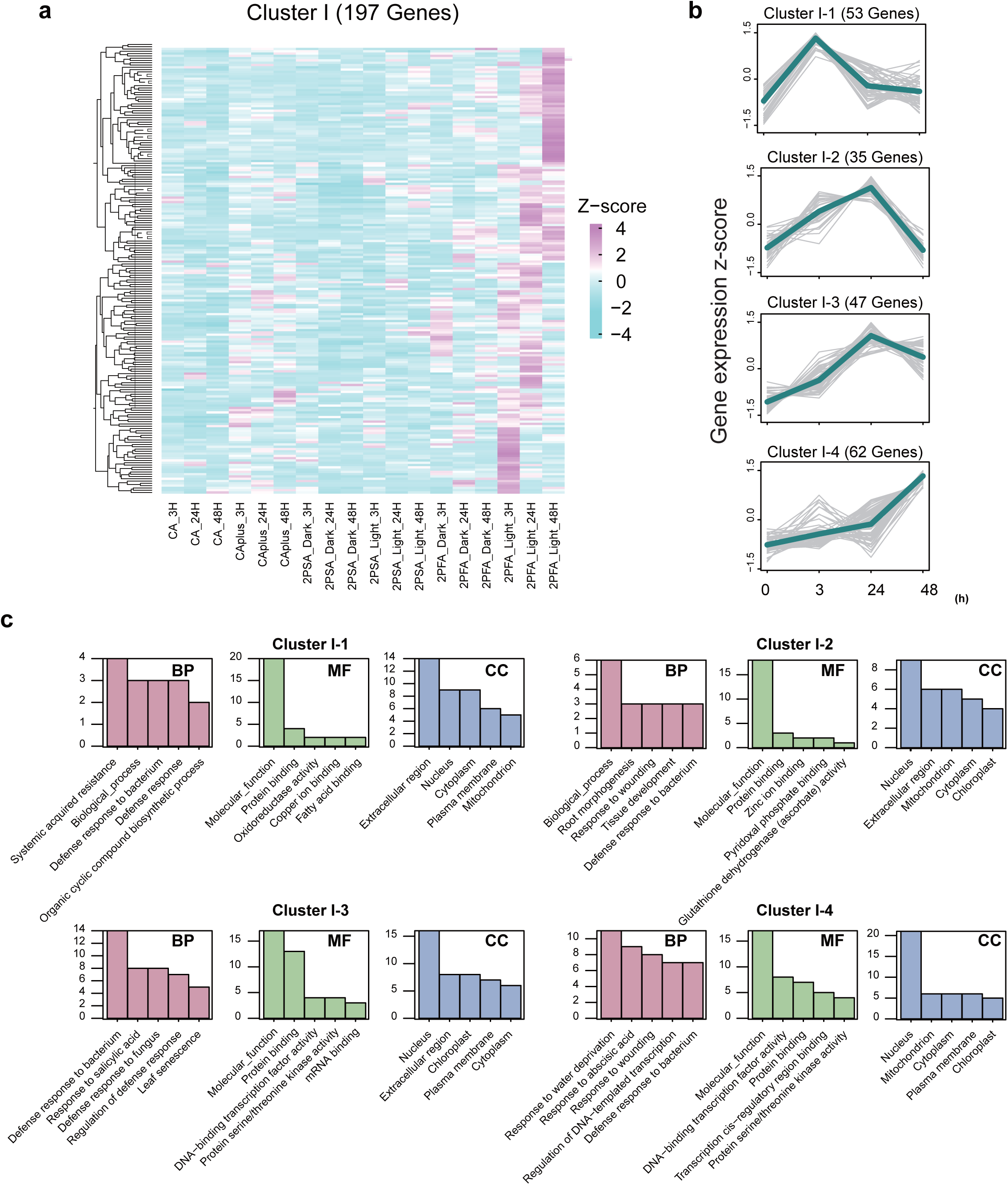
Heatmap showing expression patterns among groups (CA, CA+, 2PSA, and 2PFA) by *k-means* analysis and Gene Ontology (GO) analysis in cluster I. (a) Heatmap of time-series expression levels of genes belonging to cluster I (*n*=197) in Fig. 2b**,c** among groups (CA, CA+, 2PSA, and 2PFA). The gene expression of the time series was transformed into a Z-score. (b) Cluster analysis of 2PFA plants under light conditions in cluster I. The Z-score-transformed time-series expression levels of the genes were classified into 4 groups (Cluster I-1 to I-4) using the *k*-means method. The thin gray lines represent the individual gene expression trajectories of the genes within the cluster. A bold line represents the mean trajectory for the cluster. (c) GO enrichment analysis of genes derived from Cluster I-1-4 as shown in (b). The horizontal axis are the enriched GO terms, and the vertical axis are the numbers of the enriched genes in each cluster. The pink columns represent the biological process (BP) GO terms; the green columns represent the molecular function (MF) GO terms; the blue columns represent the cellular component (CC) GO terms.

GO enrichment analysis was conducted to determine the biological attributes of the 197 genes (**Fig. 3c**). This analysis was used to classify the genes into three GO categories: 1) biological process (BP), 2) molecular function (MF), and 3) cellular component (CC). The results showed that the BP category associated with defense such as “Defense response to bacterium”, “Defense response”, and “Response to wounding” was enriched in all the clusters. In the MF category, “Molecular function” and “Protein binding” were the top enriched GO terms in the all clusters. In the CC category, “Nucleus”, “Extracellular region”, and “Chloroplast” were the highly enriched GO terms in the cluster I-1, I-2, and I-3, but “Extracellular region” was not enriched in the cluster I-4. These genes that were upregulated in the 2PFA plants under light conditions showed that the function and location where the gene products performed their molecular function varied with the time series. Our findings in **Fig. 1** indicated that the electron transport system in the chloroplast remains active during the 2PFA process.

This led us to hypothesize that genes related to the maintenance of electron transport are involved in the 2PFA plants under light conditions. However, we did not find any terms associated with photosynthesis and detect genes directly involved in core photosynthetic processes, such as those encoding components of the light-harvesting complex.

These results suggest that the 2PFA plants under light conditions regulate the expression of genes involved in defense response rather than the genes related to photosynthesis.

### 3.4 Upregulation of genes related to sugar metabolism and reconfiguration of the photosynthetic apparatus in the 2PFA plants under light conditions

The false discovery rate (FDR) approach was used to identify the differentially expressed genes (DEGs) regulated by 2PFA and 2PSA treatments, that is, supercooling and freezing. We compared the CA+ and other treatments to detect a wide range of genes whose expressions varied with 2PSA and 2PFA treatments. A volcano plot was used to compare and reveal the DEGs in 2PFA plants under light conditions and CA+ plants (FDR < 0.05, log fold change > 1) (**Fig. 4a**). Analysis based on likelihood ratio test (LRT) revealed that there were 466 upregulated and 170 downregulated genes in the 2PSA plants under dark conditions and 642 upregulated and 94 downregulated genes in 2PSA plants under light conditions (**Fig. 4a**). Conversely, in the 2PFA treatment, there were 994 upregulated and 126 downregulated genes under dark conditions and 762 upregulated and 59 downregulated genes under light conditions (**Fig. 4a**). These results suggest that the number of upregulated genes was higher than that of downregulated genes in the 2P process.

**Fig. 4.**
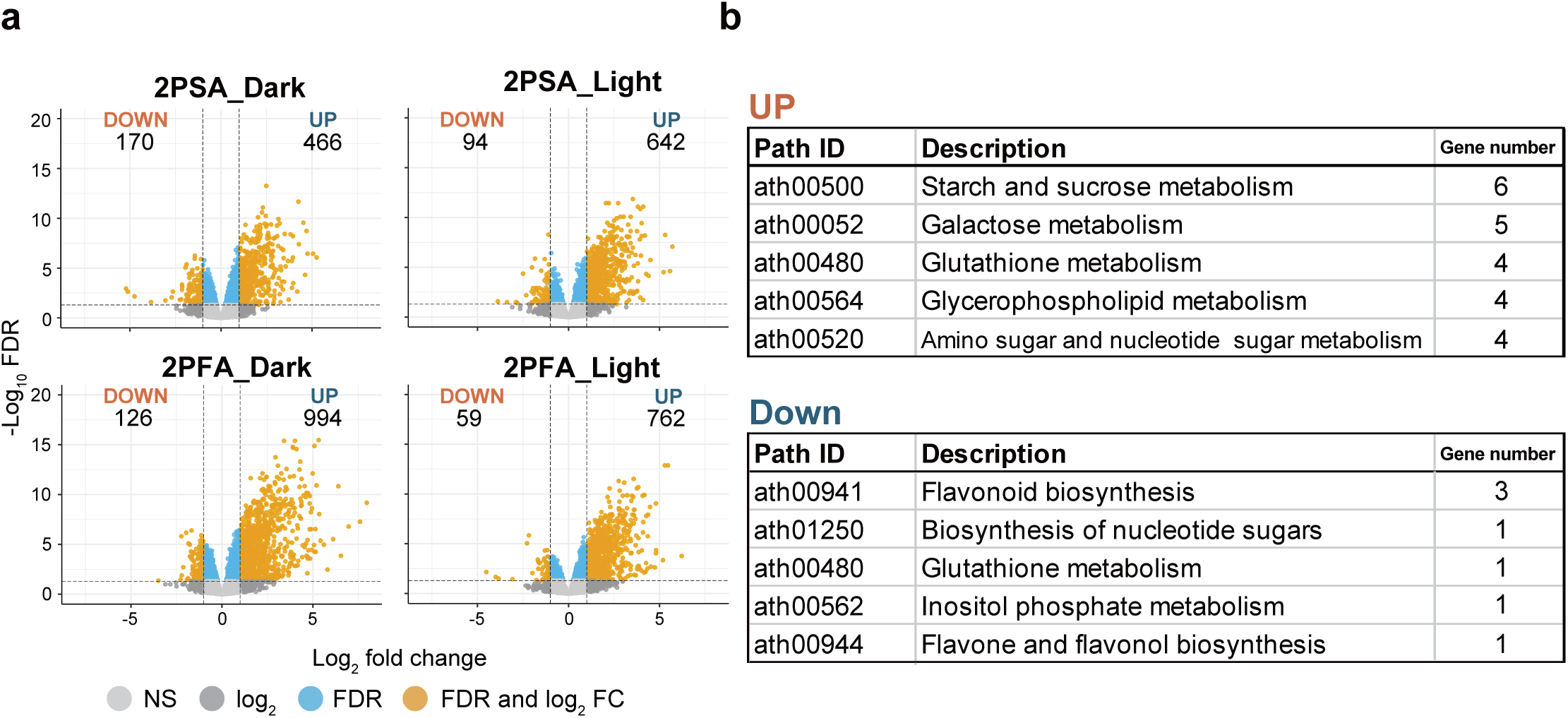
Genes related to sugar metabolisms were mainly regulated in the 2PFA plants under light conditions. (a) Volcano plots between the 2P plants (2PSA treatment or 2PFA treatment (-2°C without ice) under dark or light conditions) and CA+ plants displayed differentially expressed genes (DEGs) (the false-discovery rate (FDR) < 0.05, log2 fold-change > 1, < -1). NS (black circle) represents non-significant. The number of differentially expressed genes (DEGs) in 2P plants relative to CA+ plants. The orange columns represent the up-regulated genes; the navy blue represent the downregulated genes. (b) Kyoto Encyclopedia of Genes and Genomes (KEGG) analysis of the 821 DEGs in the 2PFA plants under light conditions relative to CA+ plants (762 upregulated, 59 downregulated genes).

To identify the key metabolic pathways during the 2PFA process under light conditions, we performed the Kyoto Encyclopedia of Genes and Genomes (KEGG) analysis. A total of 821 DEGs (762 upregulated and 59 downregulated genes) derived from 2PFA plants under light conditions were mapped onto pathways within the KEGG database (**Fig. 4b**). KEGG pathway analysis indicated that the DEGs were enriched in numerous metabolic pathways. “Starch and sucrose metabolism (ath00500)”, “Galactose metabolism (ath00052)”, “Glutathione metabolism (ath00480)”, “Glycerophospholipid metabolism (ath00564)”, and “Amino sugar and nucleotide sugar metabolism (ath00520)” were highly enriched KEGG pathways (**Fig. 4b**). As sugar metabolism was the main metabolic change in the 2PFA plants under light conditions, we visualized the expression profiles of the enriched DEGs in “Starch and sucrose metabolism” (6 genes) and “Galactose metabolism” (5 genes) as a heatmaps (**Fig. 5**). The heatmap showing the time-series expression levels of the DEGs mapped into “Starch and sucrose metabolism” revealed that the expression levels of the genes encoding the enzymes that catalyze the synthesis of trehalose, D-glucose, maltose, and sucrose, that is, *trehalose-6-phosphate synthase (TPS)*, *trehalose-6-phosphate phosphatase (TPP)*, *β-amylase (BAM)*, and *sucrose phosphate synthase 2F (SPS2F)*, respectively were upregulated in the 2PFA plants (**Fig. 5a-c**). *TPS11* and *TPPF* expressions were upregulated in 2PFA plants under the dark conditions at 24 and 48 h and light conditions at 48 h; *BAM1* expression was upregulated only in 2PFA plants under light conditions at 24 and 48 h; and *SPS2F* expression was upregulated only in CA plants at 24 and 48 h and 2PFA plants under light conditions at 48 h. In addition, a heatmap showing the time-series expression levels of the DEGs mapped into “Galactose metabolism” revealed that the expression level of the genes encoding the enzymes that catalyze the synthesis of raffinose, that is, *UDP-glucose 4-epimerase (UGE)*, *Galactinol synthase (GolS)*, and *raffinose synthase or seed imbibition protein (SIP1)* were upregulated in the 2PFA plants under light conditions (**Fig. 5d**). *UGE2* and *UGE5* expressions were upregulated in CA, 2PSA under light conditions, and 2PFA under light conditions at 24 and 48 h, suggesting that these genes were regulated in response to light. Furthermore, the expression level of *GolS2* increased only in the 2PFA plants under dark conditions at 24 and 48 h compared to that in CA, CA+, and 2PSA plants, suggesting that these genes were regulated in response to freezing.

**Fig. 5.**
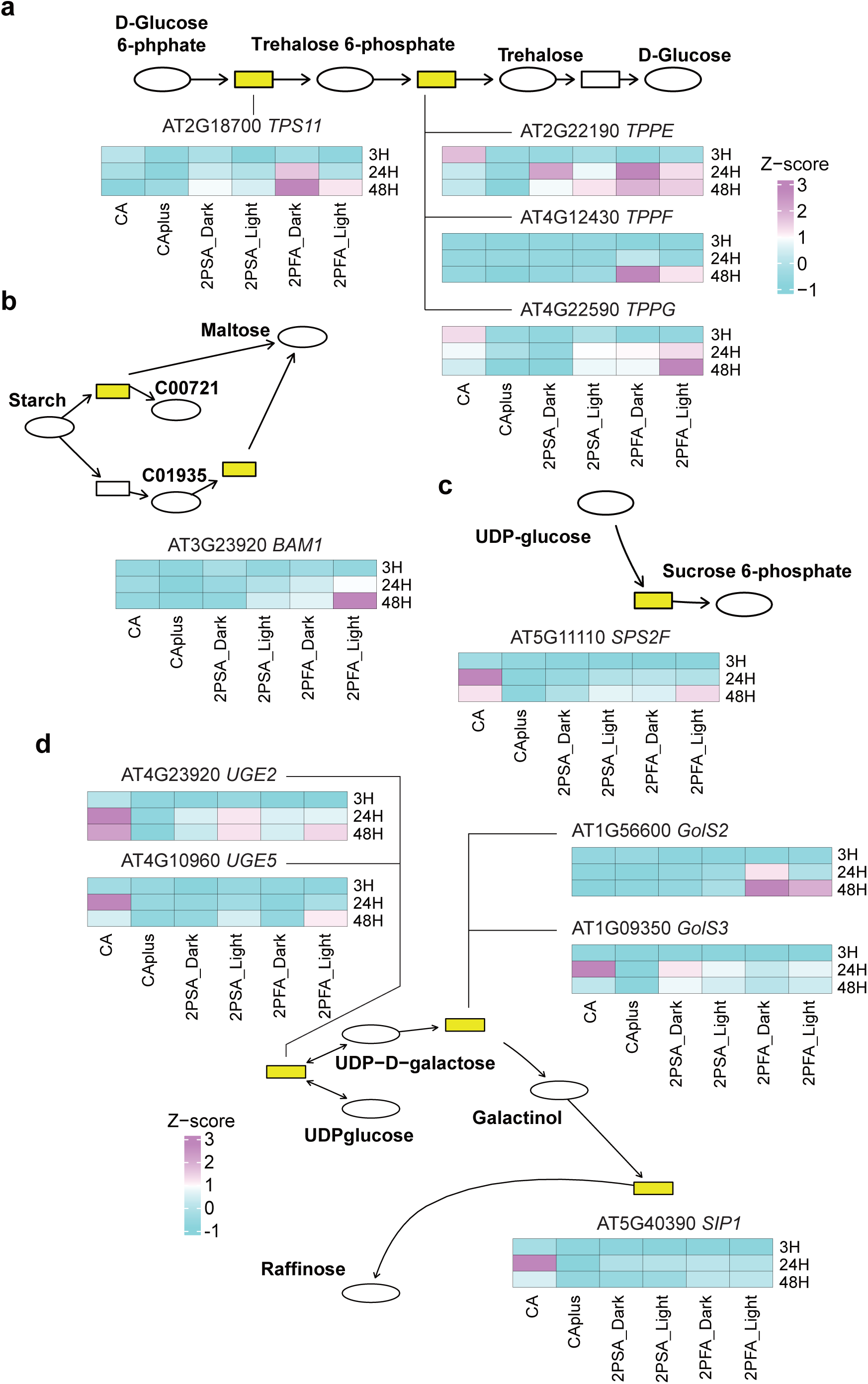
Time-series expression levels of genes enriched in “Starch and sucrose metabolism” and “Galactose metabolism”. The heatmaps showed time-series expression levels of genes enriched in “Starch and sucrose metabolism”, (a) AT2G18700; *trehalose phosphatase/synthase 11* (*TPS11*), AT2G22190; *trehalose-6-phosphate phosphatase e* (*TPPE*), AT4G12430; *trehalose-6-phosphate phosphatase f* (*TPPF*), and AT4G22590; *trehalose-6-phosphate phosphatase g* (*TPPG*), (b) AT3G23920; *beta-amylase 1* (*BAM1*), and (c) AT5G11110; s*ucrose phosphate synthase 2F* (*SPS2F*) were expressed. The heatmaps showed time-series expression levels of genes enriched in “Galactose metabolism”, (d) AT4G23920; *UDP-D-glucose/UDP-D-galactose 4-epimerase 2* (*UGE2*), AT4G10960; *UDP-D-glucose/UDP-D-galactose 4-epimerase 5* (*UGE5*), AT1G56600; *galactinol synthase 2* (*GOLS2*), AT1G09350; *galactinol synthase 3* (*GOLS3*), and AT5G40390; *raffinose synthase family protein* (*SIP1*) were expressed. All genes were expressed relative to CA_0h (CA_3, 24, 48h) and CA+_0h (CA+, 2PSA, 2PFA_3, 24, 48h). The color bar indicates gene expression transformed into Z-score for visualization.

Furthermore, GO enrichment analysis of the DEGs revealed that the most significantly enriched terms were related to general stress responses, such as “Response to water deprivation” and “Response to abscisic acid” in the 2PFA plants under light conditions (**Table S1**). To investigate the origin of sugar metabolic changes, we filtered the GO analysis results for terms associated with photosynthesis and light in the BP category (**Table S1**). Particularly, we focused on the term enriched “Photosynthesis”, “Photosynthesis, light reaction”, “Photosynthesis, light harvesting”, “Photosynthesis, light harvesting in photosystem I”, and “Photosynthesis, light harvesting in photosystem II” to detect genes associated with core photosynthetic processes directly. Among the DEGs in these categories, we particularly highlighted key genes such as *sigma factor 5 (SIG5)* and *light-harvesting chlorophyll-protein complex II subunit B1 (LHB1B1)* (**Table S2**), both of which have direct impacts on the photosynthetic machinery. SIG5 is known to maintain photosynthetic efficiency during short-term freezing by increasing PSII D2 and PSAC protein abundance (Cano-Ramirez et al., 2023). Upregulation of *SIG5* expressions was observed not only in CA at 24 and 48 h, but also in the 2PSA and 2PFA at 24h (**Table S2**). In addition, the expression of *LHB1B1*, a core component of light-harvesting chlorophyll-protein complex II (LHCII), was upregulated in 2PFA plants under light conditions at 48 h and 2PFA plants in the dark conditions at 3h, 24h, and 48h (**Table S2**).

These results suggest that changes in sugar metabolism are key metabolic pathways in 2PFA plants under light conditions. We also revealed these plants upregulated the genes related to reconfiguration of the photosynthetic apparatus. Therefore, we speculate that the accumulation of sugars plays a role in strengthening freezing tolerance induced by 2PFA treatment.

### 3.5 2PFA plants under light conditions increased the sugar concentrations

Sugars as a compatible solute can accumulate to prevent cellular dehydration and the resultant accumulation of toxic substances (Steponkus 1984; Wanner & Junttila, 1999). RNA-seq analysis (as shown in **Figs. 4,5**) revealed that the expression of genes related to sugar metabolism was regulated in 2PFA plants under light conditions. Thus, we determined their sugar concentration (**Fig. 6a, S1a**). We measured soluble sugar contents, raffinose (Raf), sucrose (Suc), glucose (Glc), and fructose (Fru). Total leaf sugars accumulated gradually within 72 h (**Fig. 6a**). Similar to the CA+ plants, 2PSA plants did not accumulate sugars under dark conditions but showed significant sugar accumulation from 24 to 72 h under light conditions. Sugar content did not change in the 2PFA plants under dark conditions until 72 h, but it clearly increased in 2PFA plants under light conditions. Under light conditions, 2PFA plants accumulated more sugar than 2PSA did..

**Fig. 6.**
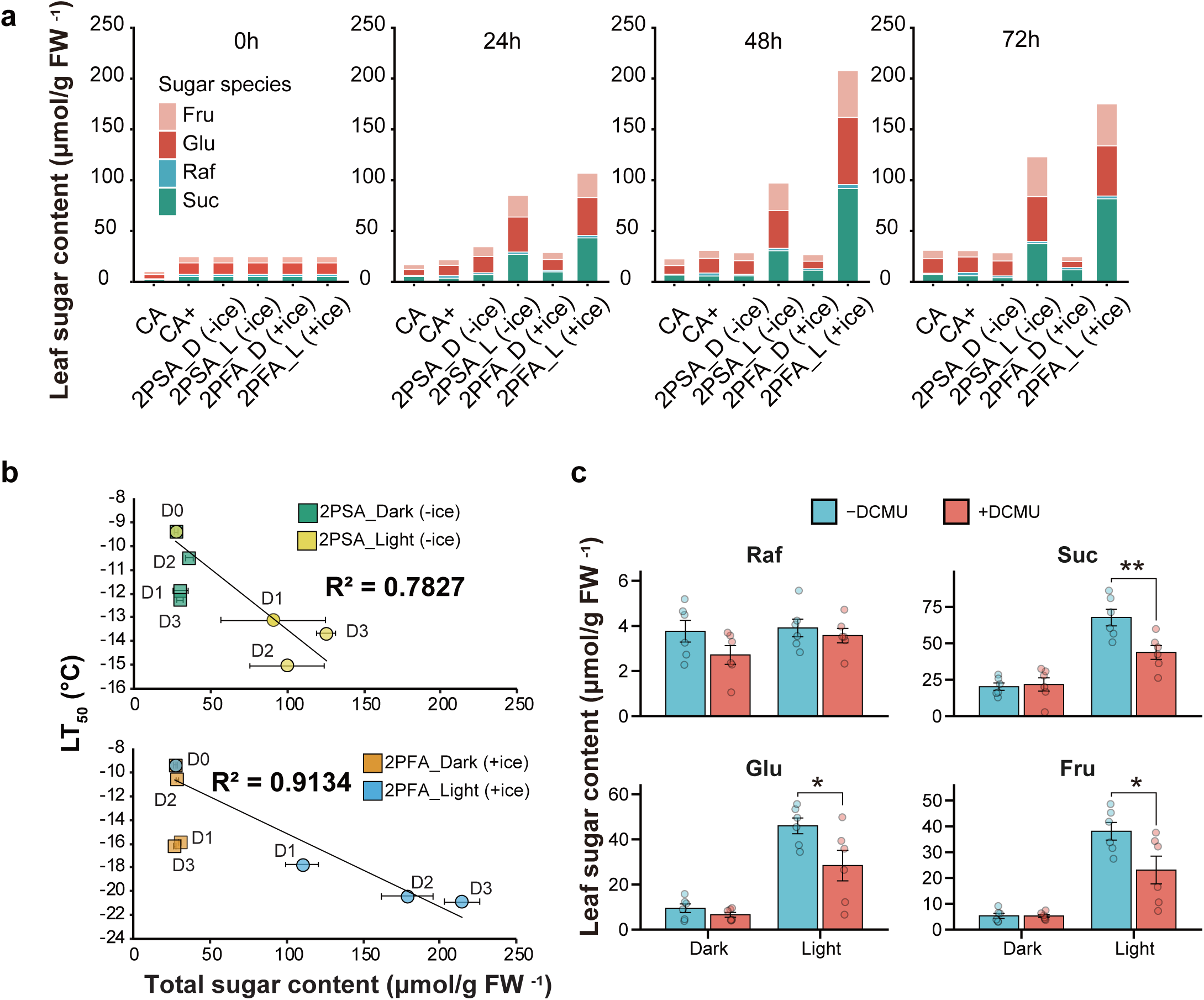
Accumulation of soluble sugars is involved in enhancement of freezing tolerance in 2PFA under light conditions via the photosynthetic electron transfer pathway. (a) Measurement of sugar concentration in the plants under various treatments (CA, CA+, 2PSA, and 2PFA). Each sugar of raffinose, sucrose, glucose, and fructose were measured in CA, CA+, 2PSA, and 2PFA plants at 0, 24, 48, and 72 h. Error bars indicate SE (*n*=3). (b) Increased sugar content strongly correlates with freezing tolerance of 2PSA and 2PFA plants under light conditions. The freezing tolerance, osmotic level, and total soluble sugar contents including raffinose, sucrose, glucose, and fructose were measured in the 2PSA and 2PFA plants at 1,2,3 days (D1, D2, D3) at - 2°C in the dark or under an 12-h light/12-h dark. The graphs indicate correlation between LT_50_ and osmotic level in 2PSA, in 2PFA plants or sugar content in 2PSA, (d) in 2PFA plants. Error bars indicate SE (*n*=3). Data for the freezing tolerance of D1, D3 in 2PFA plants and D3 in 2PSA plants were used from Sugita et al. (2024). (c) Measurement of sugar content of 2PFA plants treated with DCMU. CA plants (cold-acclimated for 7 days at 2°C) were treated with 50 μM DCMU at 2°C under green filtered low light and kept at 2°C for 1 h under dark conditions before 2PFA treatment (- 2°C with ice for 3 days under a 12-h light/12-h dark photoperiod or in the dark). Each sugar of raffinose, sucrose, glucose, and fructose was measured in the plants at 72 h after transferring to a growth chamber at -2°C. Error bars indicate SE (*n*=6). Asterisks at each temperature show significant difference (Student’s *t*-test, * *p* < 0.05, ** *p* < 0.01).

CA plants gradually accumulated Suc, Glc, and Fru for over 72 h (**Figs. 6a, S1a**). Raf slightly increased at 72 h in the CA plants. CA (CA+) after 7 d slowly accumulated Raf over time but did not accumulate other sugars. The sugar content in the 2PSA plants under the dark conditions did not change over time; however, the Suc, Glc, and Fru levels increased under light conditions, similar to those observed in the CA treatment. The sugar content in the 2PFA plant under dark conditions did not change over time; however, under light conditions Suc, Glc, and Fru slightly increased over time and Raf slightly increased at 48 h. Under light conditions, 2PFA had increased Suc content compared to 2PSA. The accumulation of these sugars was strongly and positively correlated with freezing tolerance (**Fig. 6b, S2**).

These results suggest that Suc, Glc, and Fru contents increase to enhance freezing tolerance during the 2P process under light conditions. Increased sugar content strongly correlated with freezing tolerance in 2PSA and 2PFA plants under light conditions.

### 3.6 Chloroplast redox state is involved in the 2PFA-induced increase and sugar content under light conditions

The use of DCMU in a previous study revealed that photosynthetic electron transport pathways may be involved in the enhancement of freezing tolerance in 2PFA under light conditions (Sugita et al., 2024). Therefore, we hypothesize that the accumulation of soluble sugar, which is responsible for the enhancement of freezing tolerance in 2PFA under light conditions via the photosynthetic electron transfer pathway. We measured the soluble sugar content of 2PFA plants after DCMU treatment. The plants were treated with DCMU and subsequently subjected to 2PFA treatment for 72 h under light conditions. As shown in **Fig. 6c**, DCMU pretreatment reduced the concentration of soluble sugars in 2PFA plants under light conditions at 72 h. Conversely, sugar accumulation did not change under dark conditions with or without DCMU pretreatment. DCMU pretreatment decreased Suc, Glc, and Fru concentrations in 2PFA plants under light conditions but showed no effect under dark conditions (**Fig. 6c**).

These results suggest that the photosynthetic electron transport pathway is involved in the process of 2PFA under light conditions and increases sugar accumulation.

## 4 Discussion

### 4.1 Light is essential for sugar accumulation during the 2PFA process

In the CA process of herbaceous plants, electron transfer (Tominaga et al., 2021), carbon fixation (Savitch et al., 2001) and sugar accumulation (Wanner & Junttila, 1999) occur during photosynthesis. It has been suggested that plants spontaneously stop their growth at non-freezing low temperatures and divert photosynthetic products to accumulate the sugars required to enhance freezing tolerance (Wingler, 2015), and it is unclear whether a similar phenomenon will occur under freezing conditions. We have previously demonstrated that the increase in freezing tolerance during the 2PFA process under light conditions was inhibited by the DCMU treatment (Sugita et al., 2024), suggesting that photosynthesis may be involved in enhancing freezing tolerance. To confirm this result, we constructed a new experimental system that could measure photosynthetic activity in intact frozen plants (**Fig. 1**). Using this new experimental system, we measured the *Fv/Fm* values of *Arabidopsis* under the freezing conditions at -2°C. The measurements indicate that electron transfer possibly occurred in the chloroplast during the 2PFA process (**Fig. 1e**). In this study, our RNA-seq data showed the upregulation of genes associated with reconfiguration of the photosynthetic apparatus (*SIG5* and *LHB1B1*) (**Table S2**). Given that *Fv/Fm* values are decreased under short-term freezing in *sig5* mutants compared to WT (Cano-Ramirez et al., 2023), upregulation of *SIG5* expression may be involved in maintaining electron transfer during the 2PFA process under light conditions. In addition, the expression of *LHB1B1* was upregulated in the 2PFA plants under light conditions (**Table S2**), which leads to an idea that the increased expression of *LHB1B1* is a strategy to maximize light harvesting under freezing conditions. While *k*-means and GO analysis did not identify any terms directly associated with photosynthesis in the Cluster I, which showed the upregulated expression only in the 2PFA plants (**Fig. 2**, **3**), clusters except for cluster I-1 were enriched in the “Chloroplast” of the CC category (**Fig. 3c**). We thought that this finding might be able to identify genes indirectly involved in photosynthesis, such as those related to reactive oxygen species (ROS) removal. **Table S3** shows the genes enriched “Chloroplast” of the CC category in the Cluster I-2-4. The analysis revealed several genes associated with mitigating ROS and maintaining photosynthetic components, which are crucial for protecting the photosystems. For example, in the cluster I-3, the expression of *methionine sulfoxide reductase B8* (*MSRB8*; AT4G21840), an enzyme known for repairing oxidized proteins, was upregulated, especially at 24 and 48h. Overexpression of *MSRB8* in *Arabidopsis* has been reported to confer tolerance to oxidative stress by increasing glutathione S-transferase activity, and these plants exhibited a smaller loss of chlorophyll compared to the control under H_2_O_2_ treatment (Li et al., 2012). This finding is consistent with our KEGG analysis data, which showed “Glutathione metabolism (ath00480)” was highly enriched in the 2PFA plants under light conditions (**Fig. 4**). In addition, we identified the transcriptional regulator, *WRKY53* (AT4G23810) in the cluster I-3 (**Table S3**), whose expression was also upregulated at 24 and 48h. The overexpression of a homologous gene in non-heading Chinese cabbage has been shown to not only positively regulate chlorophyll biosynthesis but also enhance cold tolerance (Chen et al., 2025). Collectively, these transcriptional changes might allow the photosynthetic apparatus to successfully avoid damage from ROS, maximize light collection, and drive electron transfer in the chloroplast (**Fig. 1**).

Interestingly, this study also revealed that the levels of sugar, such as sucrose and glucose, notably increased during the 2PFA process only under light conditions (**Fig. 6**), and the accumulation of these sugars was suppressed by DCMU pretreatment (**Fig. 6**). In addition, the sugar concentration did not change in 2PFA plants under dark conditions, suggesting that light is essential for sugar accumulation during the 2PFA process. The *Fv/Fm* values of Antarctic moss during freezing are between 0.2 and 0.4 at approximately -5°C (Lovelock et al., 1995). Barták et al. (2007) have shown via chlorophyll fluorescence measurements that Antarctic lichen species are capable of performing primary photosynthesis at -15°C. Furthermore, photosynthetic capacity such as gas exchange and PSⅡ efficiency (*Fv/Fm*) is maintained in the evergreen alpine cushion plant *Saxifraga paniculata*, even under severely freezing conditions (Hacker & Neuner, 2006). These results showed that it is quite possible that sugars are synthesized via photosynthesis even in the 2PFA process, a phenomenon that occurs under freezing.

### 4.2 Upregulation of genes involved in sugar metabolism during the 2PFA process under light conditions

Research on freezing tolerance mechanisms has shown that sugars alleviate freezing-dehydration stress (Uemura et al. 2003); therefore, investigating the genes involved in sugar increase will help to understand the mechanisms that enhance freezing tolerance. RNA-seq analysis showed that the genes involved in sugar metabolism of *TPS*, *TPP*, *BAM*, *SPS*, *UGE*, *GolS*, and *SIP* were upregulated during the 2PFA process under light conditions (**Fig. 5**), indicating that changes in metabolic pathways include trehalose, starch, sucrose, and galactose metabolisms. Significant increases in glucose, fructose, and sucrose levels were observed in this study, especially during the 2PFA process under light conditions.

Among these sugar metabolism genes, *BAM* and *SPS* have been shown not to be only upregulated during the CA process but also involved in the enhancement of freezing tolerance in transgenic plants. The *BAM3* is upregulated in CA (Kaplan & Guy, 2005). RNAi lines with reduced expression of this gene, which increased starch and decreased maltose and sucrose levels during the initial CA process, showed reduced freezing tolerance. In addition, BAM3 proteins localize to the chloroplasts and PSII is more sensitive to freezing stress in the RNAi line. In 2PFA plants under light conditions, *BAM1* but not *BAM3* was upregulated (**Fig. 5b**). BAM1 protein is also localized to the chloroplasts (Kaplan et al., 2006) and better adapted to low temperatures than BAM3 (Storm et al., 2018). Sucrose phosphate synthase (SPS) is another key enzyme for sugar accumulation during the CA process (Nägele et al., 2012), and it has been shown to increase gene expression and protein level and activity during the CA process (Strand et al., 1997, 2003; Kaplan et al., 2007). In addition, studies using overexpression or antisense suppression mutants have shown that sucrose content varies with *SPS* expression and freezing tolerance is positively correlated with sucrose content. The *SPS2F* gene, which is upregulated during the CA process, is also expressed during the 2PFA process (**Fig. 5c**, Kaplan et al., 2007).

*TPS* and *TPP*, which are involved in trehalose metabolism, and *UGE* and *GolS*, which are involved in galactinol metabolism, are also upregulated during the CA process (Fowler & Thomashow, 2002; Oono et al., 2006; Li et al., 2008; Jia et al., 2016; Takahashi et al., 2024). TPP and GolS have been shown to be involved in increasing dehydration stress tolerance (Taji et al., 2002; Lin et al., 2019). In addition, *GolS3* is a member of the CBF regulon, and its expression is induced by low temperatures and the overexpression of *CBF3/DREB1A* (Fowler & Thomashow 2002, Taji et al., 2002). It is presumed that it is difficult for plant cells to take up liquid water from outside under extracellular freezing conditions, similar to under dehydration stress. Therefore, it is possible that the enhancement of the freezing resistance in the 2PFA process occurs through the enhancement of the dehydration stress resistance. In this study, the cluster analysis using *k*-means and GO analysis revealed that “Water deprivation” in the BP categories was enriched in the cluster I-4, in which the genes were upregulated in the 2PFA plants at 48 h (**Fig. 3b,c** and **Table S1**). These results indicate that during the 2PFA process under light conditions, plants are gradually exposed to dehydration stress over time.

UDP-glucose is a substrate for both TPS and UGE (Cabib & Leloir, 1958; Barber et al., 2006). In *Arabidopsis*, UGE2, UGE4, and UGE5, which belong to the plant UGE II family based on phylogenetic relationships, are involved in the conversion of UDP-glucose to UDP-galactose, whereas UGE1 and UGE3, which belong to the plant UGE I family, is involved in the reverse reaction (Barber et al., 2006, Kotake et al., 2009, Umezawa et al., 2024). UDP-glucose is thought to be used not only for trehalose metabolism via TPS but also for galactinol metabolism via UGE because *UGE2* and *UGE5* were upregulated in the 2PFA process. Although *TPS11* and *TPPs* were upregulated in the 2PFA plants under light conditions, there was no difference in trehalose content in the plants after 7 d of CA treatment (**Fig. S1b**). This substrate is expected to decrease if UDP-glucose also flows toward galactose metabolism. In such cases, the activity of TPS must be increased to maintain a constant trehalose content. This may explain the results of this study. In contrast, the expressions of *UGE1*, *UGE3*, and *UGE4* were upregulated under dark conditions in 2PSA and 2PFA plants. Consistently, *TPS* genes, which encode enzymes that convert UDP-Glc to trehalose-6-P (Lin et al., 2019), such as *TPS1*, *TPS6*, *TPS7*, *TPS8*, *TPS9*, and *TPS10*, were upregulated during the 2PSA and 2PFA processes under dark conditions (**Fig. S3**). In addition, *TPP* genes, which encode trehalose dephosphorylation enzymes (Lin et al., 2019), were upregulated in 2PSA and 2PFA plants under light conditions (**Fig. S3**). *TPPH* was clearly upregulated in 2PSA and 2PFA plants under dark conditions but not under light conditions. *TPPA* and *TPPE* genes in the 2P treatments also showed higher expression levels than those in CA treatment. Furthermore, the gene expression of *TPPF* is specifically increased in 2PFA plants, but this change was more pronounced under dark conditions. These results suggest that trehalose-6-P and trehalose may accumulate in both 2PSA and 2PFA plants under dark conditions. In contrast, under light conditions, 2PSA and 2PFA plants may accumulate glucose to convert it into UDP-galactose.

UDP-galactose is used not only for raffinose synthesis via GolS, but also for cell wall synthesis via galactan synthase (GALS). In this study, raffinose did not accumulate during the 2PFA process (**Fig. 6a, S1a**) despite the increased gene expression of *GOLS*, which encodes the enzyme that synthesizes the precursor of raffinose, and this could also be the result of divergent substrate utilization (**Fig. 5d**). Cell wall synthesis is also important in the enhancement of freezing tolerance. Mechanical stress due to ice crystal formation causes a different injury than freeze dehydration and also causes severe cellular damage in the tolerance mechanism of which pectin components of the cell wall have been suggested to be involved (Yamazaki et al., 2008). Homogalacturonan (HG), which is the most abundant polymer of pectin, is demethylated by regulating pectin methylesterases (PME) and pectin methylesterase inhibitors (PMEI), and the biomechanical properties of the cell wall are then changed through calcium crosslinking of HG (Ridley et al., 2001, Wormit & Usadel, 2018). A previous study showed that the overexpression of *PMEIs* in *Arabidopsis* decreased freezing tolerance (Chen et al., 2018). In addition, the regulation of water loss through the fucosylation of rhamnogalacturonan II (RGII) pectin domains is involved in the enhancement of freezing tolerance (Panter et al., 2019, 2023). The expressions of *MUR1* and *MUR2*, which are involved in the fucosylation of mannose and xyloglucan, were upregulated in 2P plants under light conditions. Conversely, *MUR3*, which encodes xyloglucan galactosyltransferase, was upregulated in the 2PFA process under both dark and light conditions (**Fig. S3**).

Accumulation of β-1,4-galactan, which forms neutral side chains of RGI, played a role in increasing freezing tolerance in cold-acclimated plants through the regulation of gene expression of *GALSs* (Takahashi et al., 2024). However, because *GALS1*, *GALS2*, and *GALS3* were not regulated in the 2P process, it is believed that there were no changes in the cell wall structure due to galactan accumulation (**Fig. S3**).

### 4.3 In future

This study revealed that sugar metabolism via photosynthesis proceeds even during the 2PFA process, and the accumulated amount is significantly higher than that during cold acclimation. These results indicate that photosynthesis occurs under freezing conditions in higher plants. However, the genes involved in sugar metabolism in the 2PFA process were those that were not used in the cold acclimation process, suggesting that there may be downstream or upstream regulation during the 2PFA process. Future research should investigate how each of these genes regulates sugar metabolism or how their expressions are regulated during the 2PFA process. In addition, although glutathione metabolism was found to be associated to the 2PFA process (**Fig. 4** and **Table 1**), 2PH studies on trees in the field have shown that freezing reduces glutathione and the activation of enzymes related to glutathione metabolism (Kuroda et al., 1990). Studies on redox regulation are also necessary to better understand freeze acclimation, as it is presumed that excess energy in photosynthesis at low temperatures is handled through substrates such as glutathione (Foyer & Noctor, 2009).

## Acknowledgment

We would like to express our thanks to Professor Maya Matsunami of Iwate University for the helpful suggestion of RNA-seq analysis. We would like to express our sincere thanks to Professor Kintake Sonoike of Waseda University for a helpful suggestion. We thank Editage for editing a draft of this manuscript (https://www.editage.jp/) and their financial assistance for young researchers. Finally, we thank Ms. Yuko Suzuki, Ms. Nozomi Yokota and Ms. Michiko Saito (Iwate University) for her technical assistance.

---

**Supporting Table 1** Gene Ontology (GO) terms in the biological process (BP), cellular component (CC) category, molecular function (MF), category enriched among differentially expressed genes (DEGs) in 2^nd^-phase freezing acclimation (2PFA) plants under light conditions. DEGs were identified in 2PFA plants under light conditions relative to CA+ plants using a likelihood ratio test (FDR < 0.05, log_2_ fold-change > 1, < -1). GO enrichment analysis was performed on the 821 DEGs (762 upregulated and 59 downregulated genes), and terms associated with photosynthesis and light were extracted from the BP category.

**Supporting Table S2** Time-series expression levels of genes enriched in Go term associated with photosynthesis and light in the BP category. Expression levels of *sigma factor 5 (SIG5)* (AT5G24120) and *light-harvesting chlorophyll-protein complex ii subunit b1* (*LHB1B1)* (AT2G34430). Gene expression levels were normalized by using Fragments Per Kilobase of exon per Million mapped fragments (FPKM).

**Supporting Table S3** Time-series expression of genes enriched in “Chloroplast” in cluster I-2 to I-4 in the 2PFA plants under light conditions at 3, 24, and 48 h. The list showed the genes belonging to the cluster I-2 to I-4 as shown in Fig. 3 and determined as enriched shown in “Chloroplast” in the cellular component (CC) category. Gene expression levels were normalized by using Fragments Per Kilobase of exon per Million mapped fragments (FPKM), which is the expression level normalization method. Although “Chloroplast” was also enriched in cluster I-1, it is not shown in **Fig. 3c** because only the top five GO terms in each category are displayed. The time-series expression levels of the genes annotated with “Chloroplast” in clusters I-1 is listed in.

**Supporting Information S1** Soluble sugars in 2PFA under light conditions were accumulated that show greater than those observed in CA, CA+, and 2PSA plants. (a) Measurement of sugar concentration in the plants under various treatments (CA, CA+, 2PSA, and 2PFA). Each sugar of raffinose, sucrose, glucose, and fructose in CA, CA+, 2PSA, and 2PFA plants at 0, 24, 48, and 72 h, that showed in Fig. 6a, was plotted as a line graph. Error bars indicate SE (*n*=3). (b) Measurement of trehalose content in the 2PFA plants under light 34 conditions. Trehalose was measured in the plants at 0 and 72 h after transferring to the freezing chamber at -2°C.

**Supporting Information S2** Correlation between the LT_50_ and raffinose, sucrose, glucose, fructose in (a) 2PSA and (b) 2PFA plants under light conditions at 1,2,3 days (D1, D2, D3) at -2°C in the dark or under an 12-h light/12-h dark. Error bars indicate SE (n=3). Data for the freezing tolerance of D3 in 2PSA plants and D1, D3 in 2PFA plants were used from Sugita et al. (2024).

**Supporting Information S3** Time-series expression levels of genes related to metabolism of UDP-glucose and UDP-galactose. The heatmaps showed time-series expression from 3h to 48h. AT1G78580; *trehalose phosphatase/synthase 1* (*TPS1*), AT1G16980; *TPS5*, AT1G68020; *TPS6*, AT1G06410; *TPS7*, AT1G70290; *TPS8*, AT1G23870; *TPS9*, AT1G60140; *TPS10*, AT5G51460; *TREHALOSE-6-PHOSPHATE PHOSPHATASE A* (*TPPA*), AT2G22190; *TPPE*, AT4G12430; *TPPF*, AT4G22590; *TPPG*, AT4G39770; *TPPH*, AT5G65140; *TPPJ*, AT4G23920; *UDP-D-glucose/UDP-D-galactose 4-epimerase 2* (*UGE2*), AT4G10960; *UGE5,* AT1G12780; *UGE1*, AT1G63180; *UGE3*, AT1G64440; *UGE4*, AT3G51160; *MUR1*, AT2G03220; *MUR2*, AT2G20370; *MUR3*, AT1G30620; MUR4, AT5G15650; *MUR5*, AT3G59770; *MUR11*, AT2G47180; *GOLS1*, AT1G56600; *GOLS2*, AT1G09350; *GOLS3*, AT3G28340; *GOLS8*, AT2G33570; *GALS1*, AT5G44670; GALS2, AT4G20170; *GALS3* were expressed. All genes were expressed relative to CA_0h (CA_3, 24, 48h) and CA+_0h (CA+, 2PSA, 2PFA_3, 24, 48h). The color bar indicates gene expression transformed into Z-score for visualization.

## Reference

Andrzejowska A, Hájek J, Puhovkin A, Harańczyk H, Barták M. 2024. Freezing temperature effects on photosystem II in Antarctic lichens evaluated by chlorophyll fluorescence. Journal of Plant Physiology 294: 154192.

Barber C, Rösti J, Rawat A, Findlay K, Roberts K, Seifert GJ. 2006. Distinct properties of the five UDP-D-glucose/UDP-D-galactose 4-epimerase isoforms of *Arabidopsis thaliana*. Journal of Biological Chemistry 281: 17276–17285.

Barták M, Váczi P, Hájek J, Smykla J. 2007. Low-temperature limitation of primary photosynthetic processes in Antarctic lichens *Umbilicaria antarctica* and *Xanthoria elegans*. Polar Biology 31: 47–51.

Cabib E, Leloir LF. 1958. The biosynthesis of trehalose phosphate. Journal of Biological Chemistry 231: 259–275.

Cano-Ramirez D, Panter P, Takemura T, Fraine TS, Barros LL, Dekeya R, Barros-Galvão T, Paajanen P, Bellandi A, Batstone T et al. 2023. Low-temperature and circadian signals are integrated by the sigma factor SIG5. Nature Plants 9: 661–672.

Chen J, Chen X, Zhang Q, Zhang Y, Ou X, An L, Feng H, Zhao Z. 2018. A cold-induced pectin methyl-esterase inhibitor gene contributes negatively to freezing tolerance but positively to salt tolerance in Arabidopsis. Journal of Plant Physiology 222: 67–78.

Chen X, Gao Z, Yu Z, Ding Q, Qian X, Zhang C, Zhu C, Wang Y, Zhang C, Li Y et al. 2025. BcWRKY53 promotes chlorophyll biosynthesis and cold tolerance of non-heading Chinese cabbage under cold stress. Plant Physiology and Biochemistry 219: 109398.

Fowler S, Thomashow M. 2002. Arabidopsis transcriptome profiling indicates that multiple regulatory pathways are activated during cold acclimation in addition to the CBF cold response pathway. The Plant Cell 14: 1675–1690.

Foyer CH, Noctor G. 2009. Redox regulation in photosynthetic organisms: signaling, acclimation, and practical implications. Antioxidants & Redox Signaling 11: 861–905.

Fujikawa S, Jitsuyama Y, Kuroda K. 1999. Determination of the role of cold acclimation-induced diverse changes in plant cells from the viewpoint of avoidance of freezing injury. Journal of Plant Research 112: 237–244.

Gilmour S, Sebolt A, Salazar M, Everard J, Thomashow M. 2000. Overexpression of the arabidopsis *CBF3* transcriptional activator mimics multiple biochemical changes associated with cold acclimation. Plant Physiology 124: 1854–1865.

Guy CL, Huber JL, Huber SC. 1992. Sucrose phosphate synthase and sucrose accumulation at low temperature. Plant Physiology 100: 502–508.

Gordon-Kamm WJ, Steponkus PL. 1984. Lamellar-to-hexagonal_II_ phase transitions in the plasma membrane of isolated protoplasts after freeze-induced dehydration. Proceedings of the National Academy of Sciences of the United States of America 81: 6373–6377.

Hacker J, Neuner G. 2006. Photosynthetic capacity and PSII efficiency of the evergreen alpine cushion plant *Saxifraga paniculata* during winter at different altitudes. Arctic, Antarctic, and Alpine Research 38: 198–205.

Herman E, Rotter K, Premakumar R, Elwinger G, Bae R, Ehler-King L, Chen S, Livingston D. 2006. Additional freeze hardiness in wheat acquired by exposure to -3°C is associated with extensive physiological, morphological, and molecular changes. Journal of Experimental Botany 57: 3601–3618.

Hiraki H, Watanabe M, Uemura M, Kawamura Y. 2020. Season specificity in the cold-induced calcium signal and the volatile chemicals in the atmosphere. Physiologia Plantarum 168: 803–818.

Jia Y, Ding Y, Shi Y, Zhang X, Gong Z, Yang S. 2016. The *cbfs* triple mutants reveal the essential functions of *CBFs* in cold acclimation and allow the definition of CBF regulons in *Arabidopsis*. New Phytologist 212: 345–353.

Juurakko CL, Bredow M, Nakayama T, Imai H, Kawamura Y, diCenzo GC, Uemura M, Walker VK. 2021. The *Brachypodium distachyon* cold-acclimated plasma membrane proteome is primed for stress resistance. G3 Genes|Genomes|Genetics 11.

Kamata T, Uemura M. 2004. Solute accumulation in wheat seedlings during cold acclimation: contribution to increased freezing tolerance. Cryo Letters 25: 311–322.

Kamitani M, Kashima M, Tezuka A, Nagano JA. 2019. Lasy-Seq: a high-throughput library preparation method for RNA-Seq and its application in the analysis of plant responses to fluctuating temperatures. Scientific Reports 9: 7091.

Kaplan F, Guy CL. 2005. RNA interference of Arabidopsis beta-amylase8 prevents maltose accumulation upon cold shock and increases sensitivity of PSII photochemical efficiency to freezing stress. The Plant Journal 44: 730–743.

Kaplan F, Sung DY, Guy CL. 2006. Roles of β-amylase and starch breakdown during temperatures stress. Physiologia Plantarum 126: 120–128.

Kaplan F, Kopka J, Sung DY, Zhao W, Popp M, Porat R, Guy CL. 2007. Transcript and metabolite profiling during cold acclimation of Arabidopsis reveals an intricate relationship of cold-regulated gene expression with modifications in metabolite content. The Plant Journal 50: 967–981.

Klotke J, Kopka J, Gatzke N, Heyer AG. 2004. Impact of soluble sugar concentrations on the acquisition of freezing tolerance in accessions of *Arabidopsis thaliana* with contrasting cold adaptation – evidence for a role of raffinose in cold acclimation. Plant, Cell & Environment 27: 1395–1404.

Kotake T, Takata R, Verma R, Takaba M, Yamaguchi D, Orita T, Kaneko S, Matsuoka K, Koyama T, Reiter WD, Tsumuraya Y. 2009. Bifunctional cytosolic UDP-glucose 4-epimerases catalyse the interconversion between UDP-D-xylose and UDP-L-arabinose in plants. Biochemical Journal 424: 169–177.

Kuroda H, Sagisaka S, Chiba K. 1990. Frost induces cold acclimation and peroxide-scavenging systems coupled with the pentose phosphate cycle in apple twigs under natural conditions. Journal of the Japanese Society for Horticultural Science 59: 409–416.

Le M, Engelsberger W, Hincha D. 2008. Natural genetic variation in acclimation capacity at sub-zero temperatures after cold acclimation at 4 °C in different *Arabidopsis thaliana* accessions. Cryobiology, 57: 104–112.

Le M, Pagter M, Hincha D. 2015. Global changes in gene expression, assayed by microarray hybridization and quantitative RT-PCR, during acclimation of three *Arabidopsis thaliana* accessions to sub-zero temperatures after cold acclimation. Plant Molecular Biology 87: 1–15.

Lee JI, Yu DJ, Lee JH, Kim SJ, Lee HJ. 2013. Comparison of mid-winter cold-hardiness and soluble sugars contents in the shoots of 21 highbush blueberry (*Vaccinium corymbosum*) cultivars. The Journal of Horticultural Science and Biotechnology 88: 727–734.

Li CW, Lee SH, Chieh PS, Lin CS, Wang YC, Chan MT. 2012. Arabidopsis root-abundant cytosolic methionine sulfoxide reductase B genes *MsrB7* and *MsrB8* are involved in tolerance to oxidative stress. Plant and Cell Physiology 53: 1707–1719.

Li P, Ma S, Bohnert HJ. 2008. Coexpression characteristics of trehalose-6-phosphate phosphatase subfamily genes reveal different functions in a network context. Physiologia Plantarum 133: 544–556.

Lin Q, Yang J, Wang Q, Zhu H, Chen Z, Dao Y, Wang K. 2019. Overexpression of the trehalose-6-phosphate phosphatase family gene *AtTPPF* improves the drought tolerance of *Arabidopsis thaliana*. BMC Plant Biology 19.

Livingston D, Henson C. 1998. Apoplastic sugars, fructans, fructan exohydrolase, and invertase in winter oat: responses to second-phase cold hardening. Plant Physiology 116: 403–408.

Livingston D, Van K, Premakumar R, Tallury S, Herman, E. 2007. Using *Arabidopsis thaliana* as a model to study subzero acclimation in small grains. Cryobiology 54: 154–163.

Lovelock CE, Jackson AE, Melick DR, Seppelt RD. 1995. Reversible photoinhibition in antarctic moss during freezing and thawing. Plant Physiology 109: 955–961.

McKown R, Kuroki G, Warren G. 1996. Cold responses of *Arabidopsis* mutants impaired in freezing tolerance. Journal of Experimental Botany 47: 1919–1925.

Nagao M, Arakawa K, Takezawa D, Fujikawa S. 2008. Long- and short-term freezing induce different types of injury in *Arabidopsis thaliana* leaf cells. Planta 227: 477–489.

Nägele T, Stutz S, Hörmiller I, Heyer A. 2012. Identification of a metabolic bottleneck for cold acclimation in *Arabidopsis thaliana*. The Plant Journal 72: 102–114.

Oono Y, Seki M, Satou M, Iida K, Akiyama K, Sakurai T, Fujita M, Yamaguchi-Shinozaki K, Shinozaki K. 2006. Monitoring expression profiles of Arabidopsis genes during cold acclimation and deacclimation using DNA microarrays. Functional & Integrative Genomics 6: 212–234.

Öquist G, Huner NPA. 2003. Photosynthesis of overwintering evergreen plants. Annual Review of Plant Biology 54: 329–355.

Panter P, Kent O, Dale M, Smith S, Skipsey M, Thorlby G, Ian C, Ramsay N, Begum R, Sanhueza D et al. 2019. MUR1-mediated cell-wall fucosylation is required for freezing tolerance in *Arabidopsis thaliana*. New Phytologist 224: 1518–1531.

Panter P, Seifert J, Dale M, Pridgeon A, Hulme R, Ramsay N, Contera S, Knight H. 2023. Cell wall fucosylation in Arabidopsis influences control of leaf water loss and alters stomatal development and mechanical properties. Journal of Experimental Botany 74: 2680–2691.

Park S, Lee C, Doherty CJ, Gilmour SJ, Kim Y, Thomashow MF. 2015. Regulation of the Arabidopsis CBF regulon by a complex low-temperature regulatory network. The Plant Journal 82: 193–207.

Ridley B, O’Neill M, Mohnen D. 2001. Pectins: structure, biosynthesis, and oligogalacturonide-related signaling. Phytochemistry 57: 929–967.

Sakai A, Larcher W. 1987. Frost survival of plants. Berlin, Germany: Springer-Verlag.

Sasaki H. 1996. Changes in sugar content during cold acclimation and deacclimation of cabbage seedlings. Annals of Botany 78: 365–369.

Sasaki J, Sugita K, Kawamura Y. 2026 Involvement of SYT1-mediated mechanical stress tolerance in improved freezing tolerance during second phase freezing acclimation. Cryobiology and Cryotechnology 72: 13–16.

Sauter JJ, Wisniewski M, Witt W. 1996. Interrelationships between ultrastructure, sugar levels, and frost hardiness of ray parenchyma cells during frost acclimation and deacclimation in poplar (*Populus* × *canadensis* Moench ‹*robusta*›) wood. Journal of Plant Physiology 149: 451–461.

Savitch LV, Harney T, Huner NPA. 2000. Sucrose metabolism in spring and winter wheat in response to high irradiance, cold stress and cold acclimation. Physiologia Plantarum 108: 270–278.

Savitch LV, Barker-Åstrom J, Ivanov AG, Hurry V, Öquist G, Huner NP, Gardeström P. 2001. Cold acclimation of *Arabidopsis thaliana* results in incomplete recovery of photosynthetic capacity, associated with an increased reduction of the chloroplast stroma. Planta 214: 295–303.

Storm AR, Kohler MR, Berndsen CE, Monroe JD. 2018. Glutathionylation inhibits the catalytic activity of *Arabidopsis* β-Amylase3 but not that of paralog β-amylase1. Biochemistry 57: 711–721.

Strand Å, Foyer C, Gustafsson P, Gardeström P, Hurry V. 2003. Altering flux through the sucrose biosynthesis pathway in transgenic *Arabidopsis thaliana* modifies photosynthetic acclimation at low temperatures and the development of freezing tolerance. *Plant*, Cell & Environment 26: 523–535.

Strand Å, Hurry V, Gustafsson P, Gardeström P. 1997. Development of *Arabidopsis thaliana* leaves at low temperatures releases the suppression of photosynthesis and photosynthetic gene expression despite the accumulation of soluble carbohydrates. The Plant Journal 12: 605–614.

Steponkus PL. 1984. Role of the plasma membrane in freezing injury and cold acclimation. Annual Review of Plant Biology 35: 543–584.

Steponkus PL, Uemura M, Webb MS. 1993. A contrast of the cryostability of the plasma membrane of winter rye and spring ort - two species that widely differ in their freezing tolerance and plasma membrane lipid composition. In: Steponkus PL, ed. Advances in Low-Temperature Biology. London, UK: JAI Press, 2: 211–312.

Sugita K, Takahashi S, Uemura M, Kawamura Y. 2024. Freezing treatment under light conditions leads to a dramatic enhancement of freezing tolerance in cold-acclimated *Arabidopsis*. Plant, Cell & Environment 47: 2971–2985.

Song Y, Zhang X, Li M, Yang H, Fu D, Lv J, Ding Y, Gong Z, Shi Y, Yang S. 2021. The direct targets of CBFs: in cold stress response and beyond. Journal of Integrative Plant Biology 63: 1874–1887.

Takahashi, D, Gorka M, Erban A, Graf A, Kopka J, Zuther E, Hincha, D. 2019. Both cold and sub-zero acclimation induce cell wall modification and changes in the extracellular proteome in Arabidopsis thaliana. Scientific Reports 9: 2289.

Takahashi D, Johnson K, Hao P, Tuong T, Erban A, Sampathkumar A, Bacic A, Livingston D, Kopka J, Kuroha T et al. 2021. Cell wall modification by the xyloglucan endotransglucosylase/hydrolase XTH19 influences freezing tolerance after cold and sub-zero acclimation. Plant, Cell & Environment 44: 915–930.

Takahashi D, Soga K, Kikuchi T, Kutsuno T, Hao P, Sasaki K, Nishiyama Y, Kidokoro S, Sampathkumar A, Bacic A et al. 2024. Structural changes in cell wall pectic polymers contribute to freezing tolerance induced by cold acclimation in plants. Current Biology 34: 958–968.

Taji T, Ohsumi C, Iuchi S, Seki M, Kasuga M, Kobayashi M, Yamaguchi-Shinozaki K, Shinozaki K. 2002. Important roles of drought- and cold-inducible genes for galactinol synthase in stress tolerance in *Arabidopsis thaliana*. The Plant Journal 29: 417–426.

Thomashow M. 1999. Plant cold acclimation: Freezing tolerance genes and regulatory mechanisms. Annual Review of Plant Physiology and Plant Molecular Biology 50: 571–599.

Thomashow M. 2001. So what’s new in the field of plant cold acclimation? Lots! Plant Physiology 125: 89–93.

Tominaga Y, Suzuki K, Uemura M. Kawamura Y. 2021. In planta monitoring of cold-responsive promoter activity reveals a distinctive photoperiodic response in cold acclimation. Plant and Cell Physiology 62: 43–52.

Uemura M, Joseph RA, Steponkus PL. 1995. Cold acclimation of *Arabidopsis thaliana* (Effect on plasma membrane lipid composition and freeze-induced lesions). Plant Physiology 109: 15–30.

Uemura M, Warren G, Steponkus PL. 2003. Freezing sensitivity in the *sfr4* mutant of Arabidopsis is due to low sugar content and is manifested by loss of osmotic responsiveness. Plant Physiology 131: 1800–1807.

Umezawa A, Matsumoto M, Handa H, Nakazawa K, Miyagawa M, Seifert GJ, Takahashi D, Fushinobu S, Kotake T. 2024. Cytosolic UDP-L-arabinose synthesis by bifunctional UDP-glucose 4-epimerases in Arabidopsis. The Plant Journal 119: 508–524.

Yano R, Nakamura M, Yoneyama T, Nishida I. 2005. Starch-related *α* -glucan/water dikinase is involved in the cold-induced development of freezing tolerance in Arabidopsis. Plant Physiology 138: 837–846.

Wanner LA, Junttila O. 1999. Cold-induced freezing tolerance in Arabidopsis. Plant Physiology 120: 391–400.

Warren G, McKown R, Marin A, Teutonico R. 1996. Isolation of mutations affecting the development of freezing tolerance in *Arabidopsis thaliana* (L.) Heynh. Plant Physiology 111: 1011–1019.

Weiser, C. J. 1970. Cold resistance and injury in woody plants. Science 169: 1269–1278.

Wingler A. 2015. Comparison of signaling interactions determining annual and perennial plant growth in response to low temperature. Frontiers in Plant Science 5: 794.

Wormit A, Usadel B. 2018. The multifaceted role of pectin methylesterase inhibitors (PMEIs). International Journal of Molecular Sciences 19: 2878.

Yamazaki T, Kawamura Y, Minami A, Uemura M. 2008. Calcium-dependent freezing tolerance in *Arabidopsis* involves membrane resealing via synaptotagmin SYT1. The Plant Cell 20: 3389–3404.

Zhao C, Zhang Z, Xie S, Si T, Li Y, Zhu JK. 2016. Mutational evidence for the critical role of CBF transcription factors in cold acclimation in Arabidopsis. Plant Physiology 171: 2744–2759.

